# Quantitative assessment of morphology and sub-cellular changes in macrophages and trophoblasts during inflammation

**DOI:** 10.1101/2020.01.27.921908

**Authors:** Rajwinder Singh, Vishesh Dubey, Deanna Wolfson, Azeem Ahmad, Ankit Butola, Ganesh Acharya, Dalip Singh Mehta, Purusotam Basnet, Balpreet Singh Ahluwalia

**Affiliations:** Department of Physics and Technology, UiT The Arctic University of Norway, Tromsø 9037, Norway; Cell Biology and Biophysics Unit, European Molecular Biology Laboratory, Heidelberg, Germany; Department of Physics, Indian Institute of Technology Delhi, Hauz Khas, New Delhi 110016, India; Department of Clinical Science, Intervention and Technology Karolinska Univ. Hospital (Sweden); Women’s Health and Perinatology Research Group, Department of Clinical Medicine, UiT The Arctic University of Norway; Department of Obstetrics and Gynecology, University Hospital of North Norway, Tromsø, Norway

## Abstract

During inflammatory condition in pregnancy, the macrophages present at the feto-maternal junction release an increased amount of NO and pro-inflammatory cytokines such as TNF-*α* and INF-*γ*, which can disturb the trophoblast functions and thereby the pregnancy outcome. Measurement of the cellular and sub-cellular morphological modifications associated with inflammatory responses are important in order to quantify the extent of trophoblast dysfunction for clinical implication. With this motivation, we investigated morphological, cellular and sub-cellular changes in externally inflamed RAW264.7 (macrophage) and HTR-8/SVneo (trophoblast) using structured illumination microscopy (SIM) and quantitative phase microscopy (QPM). We monitored the production of nitric oxide (NO), changes in cell membrane and mitochondrial structure of macrophages and trophoblasts when exposed to different concentration of pro-inflammatory agents (LPS and TNF-*α*). *In vitro* NO production by LPS-induced macrophages increased 22-folds as compared to controls, whereas no significant NO production was seen after TNF-*α* challenge. Under similar conditions as with macrophages, trophoblasts did not produce NO following either LPS or TNF-*α* challenge. Super-resolution SIM imaging showed changes in the morphology of mitochondria and plasma membrane in macrophages following LPS challenge and in trophoblasts following TNF-*α* challenge. Label-free QPM showed a decrease in the optical thickness of the LPS-challenged macrophages while TNF-*α* having no effect. The vice-versa is observed for the trophoblasts. We further exploited machine learning approaches on QPM dataset to detect and to classify the inflammation with an accuracy of 99.9% for LPS-challenged macrophages and 98.3% for TNF-*α*-challenged trophoblasts. We believe that the multi-modal advanced microscopy methodologies coupled with machine learning approach could be an alternative way for early detection of pregnancy related inflammation after clinical studies.

## 1. Introduction

Macrophages play an important role in immune response, tissue development, remodelling and repair[1, 2]. They are generally classified into two categories depending on how they respond to the various environmental signals: classically activated macrophages (M1) and alternatively activated macrophages (M2). M1 macrophages are stimulated by the virus infections, endotoxin (LPS) or some cytokines e.g. TNF-*α*, INF-*γ*, etc. Important cytokines, such as IL-1, IL-6, IL-12 and TNF-*α* are released in response. M2 macrophages help in the tissue remodelling and repair and are characterised by the release of the cytokines such as IL-2*β* and IL-10[3, 4]. In a pregnant woman placental decidua contains 20-30 % macrophages of the total population of the leukocytes. During peri-implantation period, the decidual macrophages are inclined towards M1 phenotype. Their profile predominantly shifts towards M2 macrophage phenotypes during the pregnancy. Macrophages play important role in the spiral artery remodelling and the trophoblast invasion by clearing the apoptotic cells in the decidua[5, 6].

Better communication between the fetal trophoblast and maternal immune cells is very important for the successful outcome of a pregnancy. The trophoblast, just like an innate immune cell, expresses pattern recognition receptors (PRR) that act as ‘sensors’ of the surrounding environment[7]. Through PRR, the trophoblast can recognize the presence of pathogens, dying cells and damaged tissue[8]. Upon recognition, the trophoblast will secrete certain cytokines that in turn, will act upon the immune cells within the decidua (i.e. macrophages, T regulatory cells, NK cells), recruiting and educating them to work together in support of the growing fetus[7-9]. A viral or bacterial infection may perturb the harmony of the cross-talk between macrophages and trophoblasts which might lead to various pregnancy complications[10]. One of the major pathogens causing these infections is gram negative bacteria. These bacteria colonise the genitourinary tract of women, where they continuously release an endotoxin called lipopolysaccharide (LPS). LPS is present on the outer membrane of the gram-negative bacteria which induces inflammation by stimulating the immune system, particularly macrophages[11]. Classically activated macrophages produce TNF-*α* and nitric oxide (NO) in abundance which has been linked with pre-eclampsia, preterm delivery and early abortion[12, 13].

Several studies have been conducted to understand the mechanisms of inflammation in macrophages and trophoblasts following stimulation with various cytokines. However, we have insufficient information about the effect of LPS and other cytokines released in its effect on the morphology of these cells at the sub-cellular level. Mitochondria produces reactive oxygen species (ROS) continuously during respiration[14]. In pathological state ROS can be overproduced and thus can cause oxidative stress (OS)[15]. OS can lead to mitochondrial swelling and initiate an apoptotic cascade[16, 17]. Superoxide radical (O_2_^.-^) may also react with NO produced during infection to produce a toxic substance peroxynitrite (ONOO^-^) damaging the cells[18]. There have been few studies carried out using electron microscopy which suggest that the mitochondrial morphology of trophoblasts is altered under pathological conditions[19, 20], but these studies are limited to fixed cell due to incompatibility of electron microscopy with live cell imaging. So far to the best of our knowledge studies have not been done on live macrophages and trophoblasts using super-resolution imaging. Plasma membrane also play an important role during inflammation. PRR are generally expressed on the plasma membrane and after recognising any foreign molecule, signalling cascade is initialised which instructs a cell to produce cytokines. Therefore, the study of plasma membrane and mitochondria is crucial to mark the changes during inflammation. Many important details in the inflammation-related sub-cellular processes in these cells could have not been observed due to the limited spatial resolution of conventional fluorescence microscopy systems. Moreover, multi-modal imaging complemented with the chemical analysis are required to obtain better understanding of the inflammation related changes in macrophages and trophoblasts.

Structured illumination microscopy (SIM) is a wide-field super resolution optical microscopy technique having the resolution twice compared with the conventional microscopes[21]. Among the existing super-resolution optical microscopy techniques, SIM offers advantage of relatively high-speed, threedimensional imaging and most importantly compatible for the live cell imaging[21, 22]. Additionally, SIM is also compatible with wide range of conventionally used bright and stable fluorophores making it popular choice for imaging and optical sectioning of live biological cells[22, 23]. SIM is fluorescence-based technique and therefore requires exogenous labelling agent for imaging cells. In label-free optical imaging, quantitative phase microscopy (QPM) is a popular technique for the quantitative analysis of live biological specimens[24]. QPM techniques are capable to quantify various parameters associated with biological specimen, such as cell dynamics (fluctuations in cell thickness and/or RI), cell morphology and cell dry mass (non-aqueous content)[24-27]. Digital holographic microscopy (DHM) is one of the interferometric QPM techniques to extract the quantitative information of the live cell[26]. It is very useful technique for the single cell analysis with ease of operation and compatibility[27, 28]. In connection with vigorous algorithms of numerical reconstruction of interferograms for data measurement, DHM provides a fascinating window in modern microscopy for quantitative analysis of the specimen. In addition, single shot DHM techniques can be used for the measurement of dynamic fluctuation of the live cells[29]. The 3D cell shape can be obtained from quantitative phase which having information about the integral cell refractive index and the cell thickness[26]. This enables to determine various parameters of the cells such as volume, surface area, sphericity, etc[26, 30], which could be studied as bio-marker of cell inflammation.

In this work, we have applied different optical modalities for the observation of various morphological changes on cellular and sub-cellular levels in macrophages and trophoblasts after inducing inflammation with LPS or TNF-*α* at different concentrations. We observed the effects of various concentrations of LPS or TNF-*α* on NO production as well as cell membrane and mitochondria of macrophages and trophoblasts at different time intervals. The results suggest that these techniques together provide the clear visual and quantitative insights about the morphological and sub-cellular level changes in macrophages and trophoblasts at normal and pathological conditions. The basic information obtained in present study could be useful for the treatment strategy in the future for the cases of the infection and inflammation in pregnancy.

## 2. Materials and Methods

### 2.1 Materials

RPMI (Roswell Park Memorial Institute) 1640 medium (#R8758), LPS (#L2880), TNF-*α* (#T5944), sulphanilamide (#S9251), naphthalene diamine dihydrochloride (#N91250) and Dulbecco’s PBS (Phosphate-buffered saline #D8537) were bought from Sigma-Aldrich, St Louis, MO, USA. Agilent 8453 UV-Visible Spectrophotometer, Agilent Technologies, Santa Clara, USA. FALCON 24-Well cell culture plates, Corning Incorporated, New York, USA. Zeiss standard binocular microscope, Carl Zeiss, West Germany with 10X and 40X objective lenses. CellMask Green (CMG) (#C37608), MitoTracker Green (MTG) (#M7514) and live cell imaging solution (#A14291DJ) were bought from Thermo Fisher Scientific (Waltham, USA). #1.5, 25 mm round coverslips (#631-0172) were bought from VWR international. Human fibronectin purified from human plasma by affinity chromatography on Gelatin Sepharose 4B was a kind gift from Vascular Biology Research Group, Department of Clinical Medicine, UiT-The Arctic University of Norway. Commercial OMX 3D-SIM v4 blaze system (GE Healthcare, USA) and custom build quantitative phase microscopy system are used for SIM and QPM, respectively, at Optical Nanoscopy Lab, UiT-The Arctic University of Norway.

### 2.2 Cell culture

Macrophages (RAW 264.7) and trophoblasts (HTR-8/SVneo) cell lines used in the experiments were bought from ATCC. Cell culture was carried out in laboratory of Women’s Health and Perinatology Research group, UiT-The Arctic University of Norway. Both cell lines were cultured separately in a humidified atmosphere of 95% air and 5% CO_2_ at 37 ^o^C with glutamine containing RPMI-1640 medium supplemented with 10% fetal bovine serum and antibiotics (penicillin and streptomycin). The cells were subcultured every 2-3 days by transferring the cells to the next passage and were utilised for experiments at 80-90% confluency.

For QPM measurements, the cells were seeded in PDMS chambers located on reflecting silicon slides. Samples was incubated for 24 h and later cells were divided into two batches (a) control and (b) challenged with inflammatory agents for the experiment.

### 2.3 Measurement of NO production

For NO measurements, 1 × 10^5^ cells/ml were transferred to each well of the 24-well culture plates. The cells could stabilize and adhere for 24 h in an incubator at 37^°^ C and 5% CO_2_. The cells were then treated with three different concentrations of LPS (0.1 μg/ml, 1 μg/ml, 10 μg/ml). The production of NO released in the medium were measured spectrophotometrically in terms of nitrite formation by Griess reagent[31, 32] at 540 nm using NaNO_2_ as the standard. The measurements were performed in triplets at different time points as the result shown in Fig. 1. Similar plan was adopted to measure the NO after TNF-*α* challenge at three different concentrations (10 pg/ml, 100 pg/ml and 1000 pg/ml).

**Fig.1.**
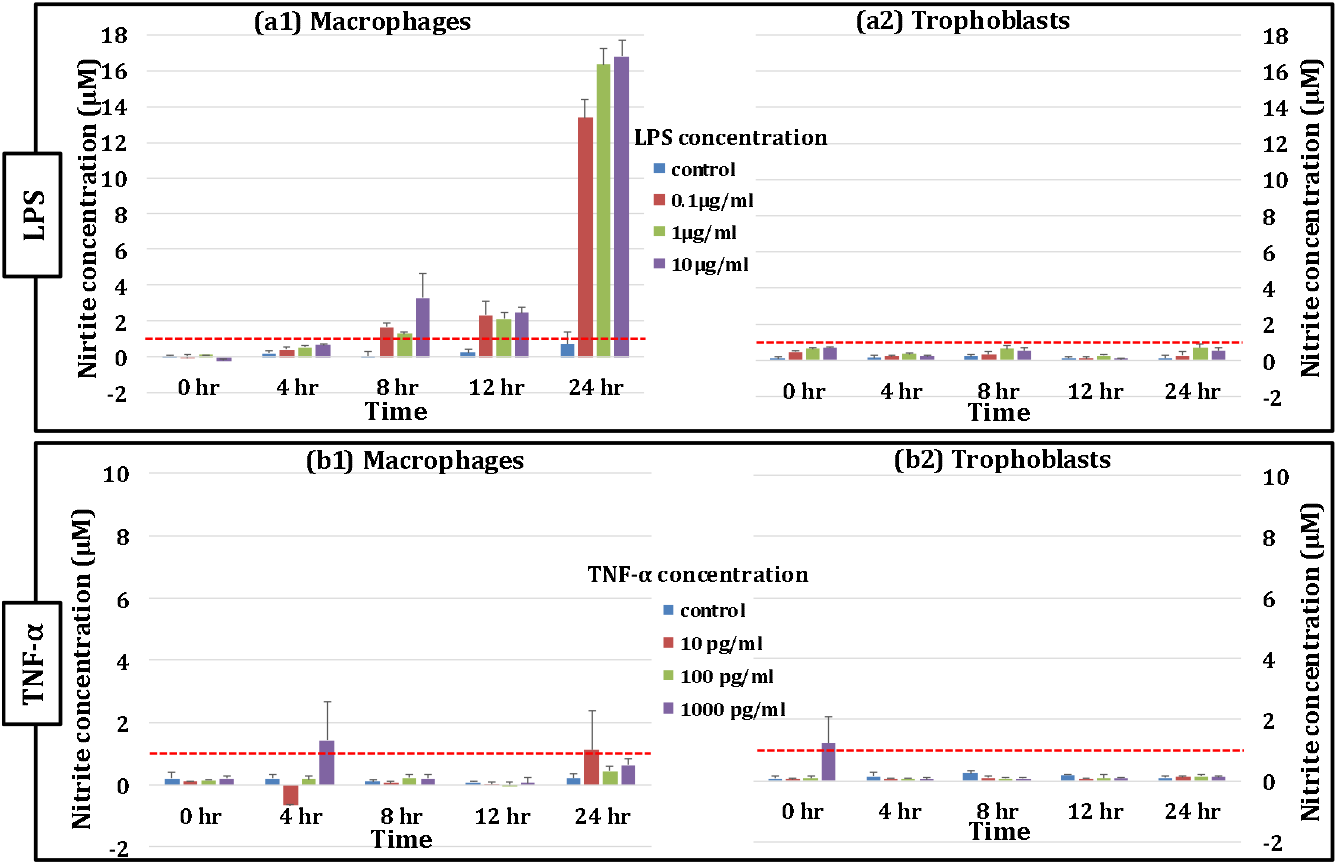
The production of NO by macrophages and trophoblasts after challenging with two inflammatory agents. (a1) macrophages and (a2) trophoblasts challenged with various concentrations LPS, (b1) macrophages and (b2) trophoblasts following TNF-α challenge at various concentrations. Bar graph shows the mean value (+S.D.) of three experiments. Considerable amount of NO was produced only in case of macrophages stimulated by LPS. The horizontal red line shows the limit of significant amount of NO production.

### 2.4 Labelling and super resolution imaging of macrophages and trophoblasts

We optimised the labelling protocol for staining cell membrane and mitochondria for both cell types using CellMask Green (CMG) and MitoTracker Green (MTG), respectively. We found that the concentration 1:1000 of CMG and 60 nM of MTG is optimum for good reconstruction of SIM images of for the both cell types. Both dyes are excited by the same wavelength (λ_ex_ = 488 nm). For labelling cell membrane, live cells on the coverslips were washed once with RPMI and then CMG (1:1000) is applied and sample is incubated for 10 min. For mitochondria labelling, live cell sample is washed once with RPMI and then incubated with MTG (60 nM) for 30 min. Then the sample is washed with RPMI thrice and mounted in imaging solution. For SIM imaging, cells were seeded on #1.5 coverslips (170 μm thick) the day before imaging and were incubated. Next day cells were challenged with an inflammatory agent and data were acquired at 2 h, 4 h, 6 h and 24 h time points. Cells were imaged at these time-points on SIM afterwards.

3D-SIM images of macrophages and trophoblasts were acquired at room temperature (T = 23 ^o^C) and within 1 h after taking out of the incubator on a commercial structured illumination microscope (DeltaVision|OMXv4.0 BLAZE, GE Healthcare). All the images were acquired using 488 nm laser with power set to either 10% or 31.5%. The images shown in this study are the projected image of z-section stacks. Each z-section is 0.125 μm thick and the thickness of samples imaged was kept between 1.2 - 2.5 μm. Exposure time was generally kept between 10 - 40 ms. After acquisition, images were reconstructed using the SoftWorX package from GE Healthcare.

### 2.5 Quantitative phase imaging of macrophages and trophoblasts

A Linnik type DHM setup is used for the imaging of the biological samples[26, 28]. Unlike to SIM microscopy, this technique does not require any labelling agent to quantify the morphological features[33]. The details of the set up can be found elsewhere [24, 26, 27]. A 60 × microscope objective (NA = 1.2) is used for the imaging of the control and inflamed live cells. He-Ne laser @632.8 nm is used to illuminate the object and record the off-axis holograms with help of CMOS camera (Hamamatsu ORCA-Flash4.0 LT, C11440-42U). These images are used further for the post processing. Fourier transform analysis is used to extract the phase delay induced by the specimen, which represents the average refractive index and thickness of specimen at all spatial points[24, 26]. The experiment is repeated several times on each set of the sample.

## 3. Results and Discussion

### 3.1 NO production measurements

The response of macrophages and trophoblasts on NO production were studied after challenging them with various concentrations (0.1 μg/ml, 1 μg/ml, 10 μg/ml) of LPS and at different time points (0 hr, 4 hr, 8 hr, 12 hr, 24 hr). After inflammation induced by LPS, the macrophages produce a large amount of NO after 24 hr as shown in Fig. 1 (a1). It can be observed from Fig. 1(a1) that the challenged cells already started producing NO compared with control, but they were below the detection limit so the results up to 4 hr are not conclusive. After 8 hr, the macrophages produce a significant amount of NO to be detected. The maximum increase in the NO production is seen between 12 - 24 hr. An increase in the NO production was expected and is in accordance with the previous studies[34, 35]. In contrast, trophoblasts did not produce any significant amount of NO even after 24 hr following LPS challenge as shown in fig 1(a2). Further, macrophages and trophoblasts were challenged with different concentrations of TNF-*α* (10 pg/ml, 100 pg/ml and 1000 pg/ml) and measured the corresponding amount of NO production in the cell culture media. Both macrophages and trophoblasts did not produce any significant amount of NO production even after 24 hr of stimulation as shown in Fig 1(b1) - (b2). Some values are above the significant amount of NO production, but the standard deviations for these readings are also large. Some negative values in the results might be due to the blank error.

In Table 1, we have listed the production of NO by macrophages and trophoblasts after inducing by the inflammatory reagents i.e. LPS and TNF-α. Only the macrophages produced NO after LPS challenge after 8 hr of inflammation and remained maximum after 24 hr. We could not measure any significant amount of increased NO production from trophoblast after LPS treatment. TNF-α could not induce the NO production to macrophages as well as trophoblasts. The red boxes of the table show the time point of the inflammation where the significant amount of NO is produced by cells and green boxes of the table showing the no significant level of NO production.

**Table 1.**
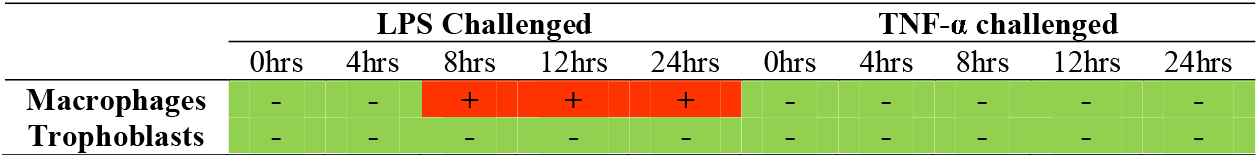
Nitric oxide measurement results obtained from Griess method for LPS and TNF-α challenged macrophages and trophoblasts for different time points.

### 3.2 SIM imaging of cell membrane and mitochondria

To detect the changes at cellular and sub-cellular level during inflammation in macrophages and trophoblasts, SIM microscopy is performed to obtain the super-resolved images of cell membrane and mitochondria. Initially, both the macrophages and trophoblasts were challenged with LPS (1 μg/ml) and TNF-*α* (1000 pg/ml) for 24 hr. The plasma membrane and mitochondria were labelled with Cell Mask Green (CMG) and MitoTracker Green (MTG), respectively. Both dyes are excited by the same wavelength (λ = 488 nm). Cell Mask, is an amphipathic molecule containing a lipophilic part for membrane loading and a negatively charged dye for membrane anchoring. The plasma membrane of the macrophages and the trophoblasts were imaged with SIM microscopy. The results show a drastic change in cell membrane morphology of LPS-challenged macrophages and TNF-*α*-challenged trophoblasts compared to the control cells as shown in Fig. 2. Figure 2(a) shows the control macrophages with defined boundaries and villi-like projections on the plasma membrane while Fig 2(b) show the spreading and the membrane disintegration after inflammation. Figure 2 (c)-(d) show the images of plasma membrane of control and TNF-*α* challenged trophoblasts. After inflammation, the cell membrane boundaries are disintegrated and the filopodia are less in numbers as compared to controls.

**Fig.2.**
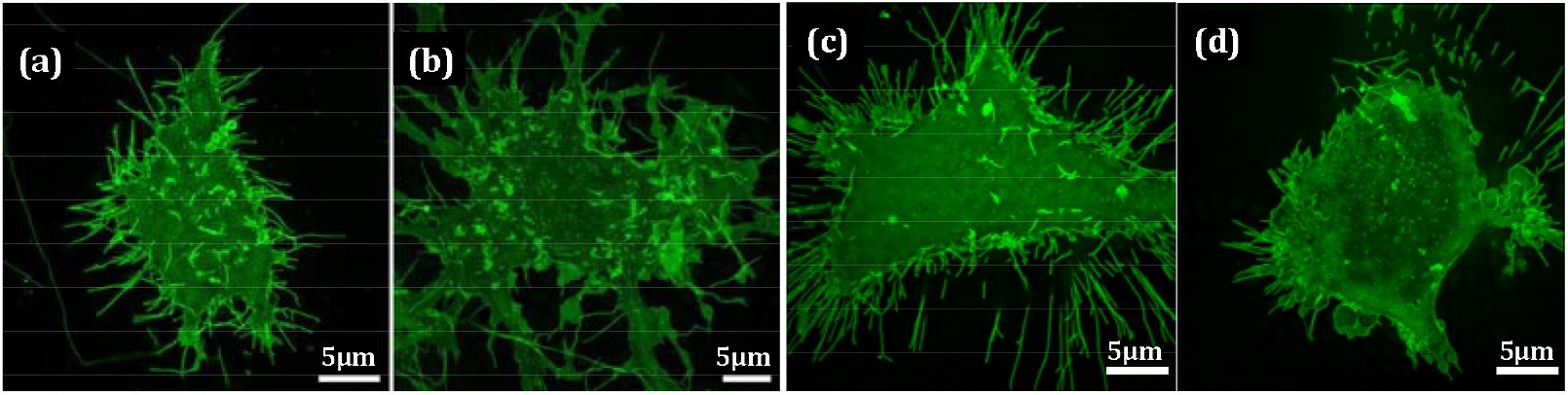
3-D projected images of plasma membrane labelled with CMG (λex= 488 nm) of (a) control and (b) LPS-challenged macrophages. Similar results obtained from (c) control and (d) TNF-α challenged trophoblasts. Comparing (a) vs. (b) and (c) vs. (d), it is observed clearly that the cell membrane integrity is lost with reduced number of filopodia after inflammation.

In addition to the cell membrane which is the first point of contact to inflammatory agents, mitochondria are another important sub-cellular organelle responsible for generating energy and thus well-being for the cell. The small size of mitochondria limits its visualization using diffraction limited optical microscope. Hence, super-resolution SIM imaging can reveal the changes in the morphology of mitochondria, especially in a live cell. Macrophages stimulated with LPS show different mitochondrial morphology as compared to the control cells. Figure 3 depicts the comparison between the mitochondrial morphology of control, LPS-challenged and TNF-*α* challenged macrophages. Fig. 3(b) suggest that the mitochondria appear to be fragmented, round and shorter in LPS-challenged macrophages compared with control. The SIM imaging show that around 50 - 60% of the cells were found with altered mitochondrial morphology after LPS-challenge (Fig. 3b). Further, Fig. 3(d)-(f) show the magnified images of the mitochondria of selected region of Fig. 3(a)-(c). Contrary, it was found that TNF-*α* challenge does not produce any remarkable change in the morphology of the mitochondria comparing to control.

**Fig.3.**
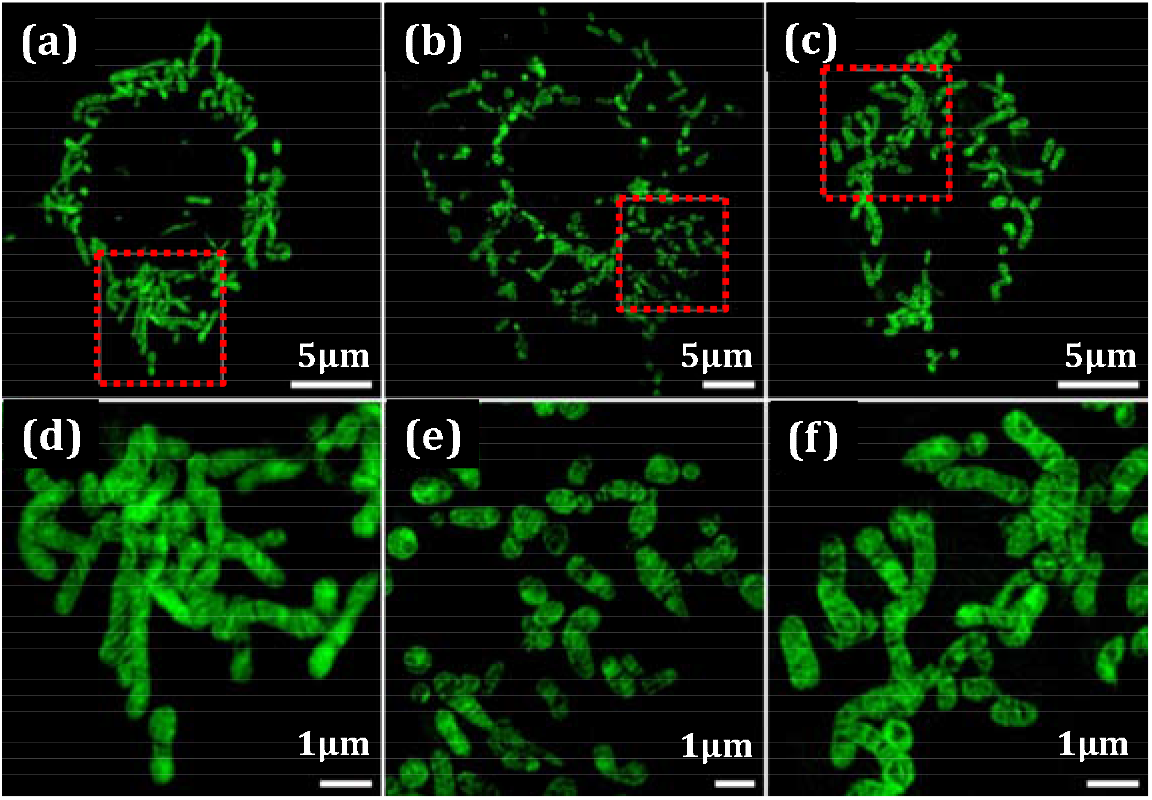
3-D projected SIM images of the mitochondria labelled with MTG (λex = 488 nm) in (a)-(c) control, LPS-challenged and TNF-α challenged macrophages, respectively. (d)-(f) are mitochondrial images of the region marked with red dotted box in (a)-(c). Mitochondria in LPS-challenged macrophages are shorter and rounded as compared to control and TNF-α challenge.

Similarly, the mitochondria of the LPS or TNF-*α*-challenged trophoblasts are also imaged and some representative images are provided in Fig. 4. Here, we did not observe significant changes on the morphology of mitochondria while inducing trophoblasts by LPS as compared to control (Figs. 4 (a) and (b)). While the morphology of the mitochondria in trophoblasts changes on treating with TNF-α as compared to control. Fig. 4(d)-(f) show the magnified images of the mitochondria of selected region of Fig. 4(a)-(c), respectively.

**Fig.4.**
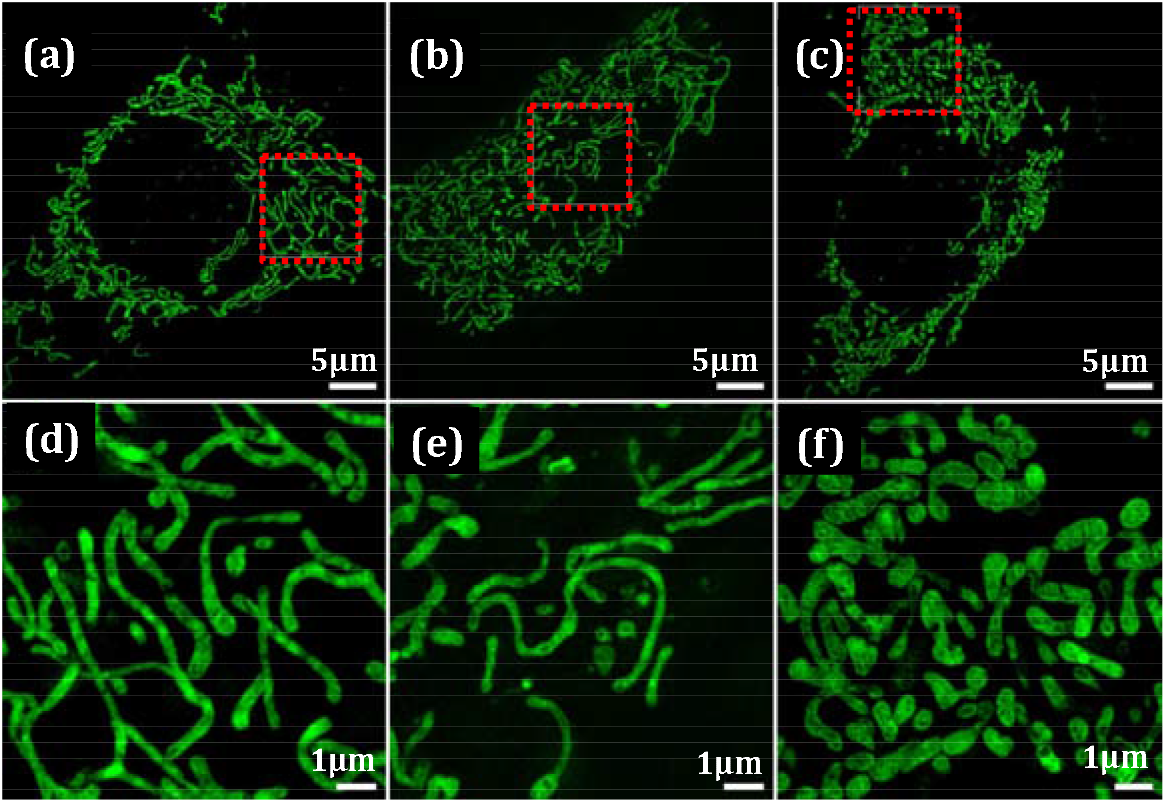
3-D projected SIM images of the mitochondria labelled with MTG (λex = 488 nm) in (a)-(c) control, LPS-challenged and TNF-*α* challenged trophoblasts, respectively. (d)-(f) are the region marked with red dotted box in (a)-(c). Mitochondria in TNF-α challenged trophoblasts are having different morphology as compared to control and LPS.

Further, we also performed quantitative analysis of the mitochondrial images of cells under different levels of treatments as described above. Image analysis was done using MiNA Single Image macro[36] as shown in Fig. 5. The features are outlined and then further analysed to get quantitative information. We have quantified number of mitochondria and their footprint area using MiNA for control and inflamed macrophages and trophoblasts. Overall, MiNA captured the visually apparent changes in the mitochondrial morphology in macrophages and trophoblasts after challenging them with LPS and TNF-α. Figure 5 (a), (c) show the mitochondria of control and LPS-challenged macrophages, respectively. MiNA detects the footprints of each individual mitochondria of Fig. 5 (a) and (c) as shown in Fig. 5 (b), (d), respectively. The footprint of mitochondria also provides the quantitative information about the projected area of the mitochondria as well as their counts.

**Fig.5.**
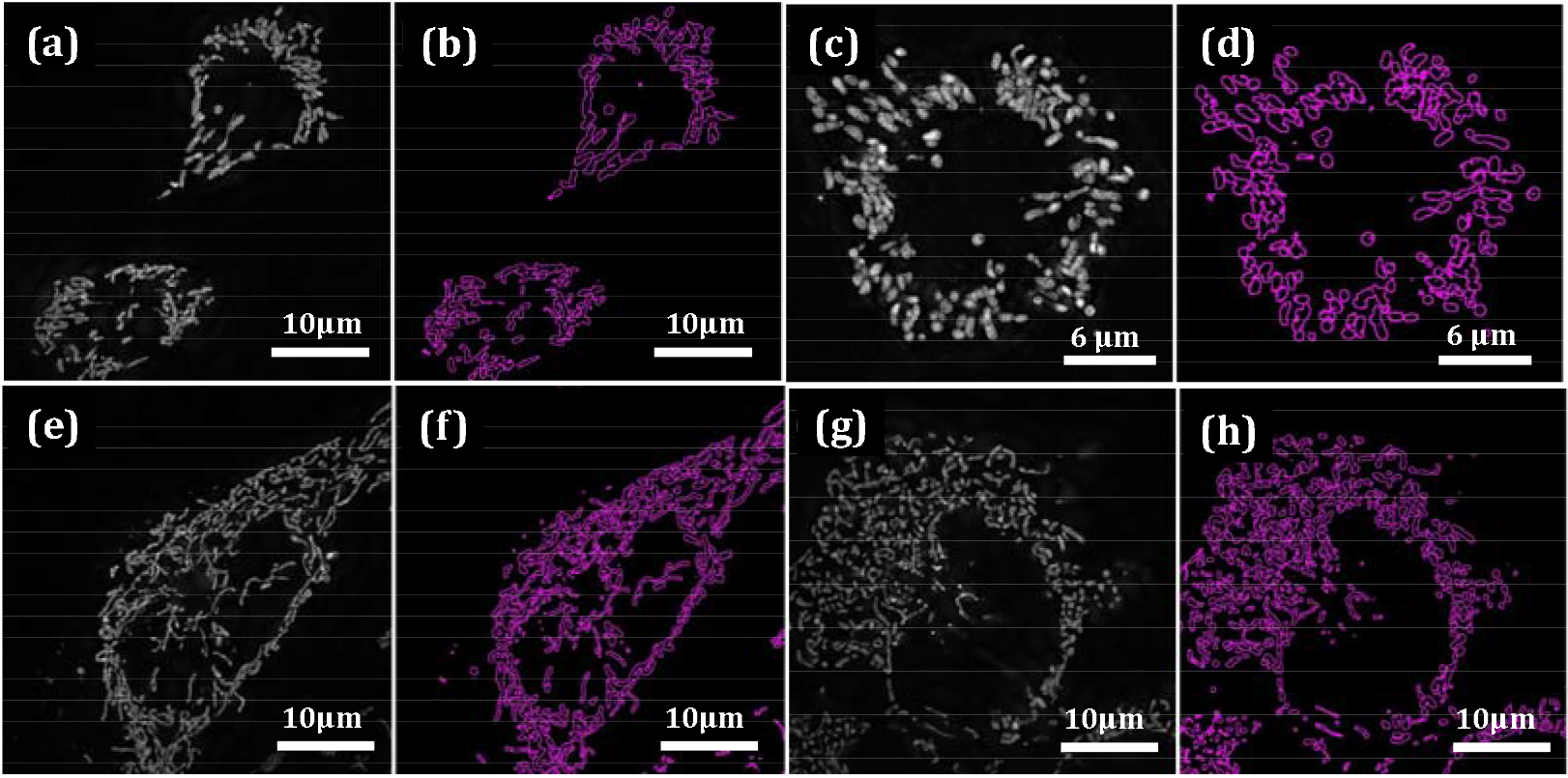
(a), (c) 3-D projected SIM images of the mitochondria in control and LPS-challenged macrophages, respectively. (b) and (d) are mitochondrial footprint corresponding to each individual mitochondria drawn by MiNA. Similarly, (e),(g) shows the 3-D projected SIM images of the mitochondria in control and TNFα challenged trophoblasts and their corresponding footprints in (f) and (h), respectively.

Figure 6 depicts the quantified parameters of mitochondria of control and inflamed macrophages and trophoblasts using MiNA. A total of 36, 33 and 40 cells for control, LPS challenge and TNF-α challenge macrophages, respectively were used in Fig. 6 (a) which shows that there is an increase in mitochondria counts in LPS challenged macrophages while TNF-*α* does not affect it. Interestingly the footprint of the mitochondria is also increased after LPS challenge macrophages while TNF-*α* challenge does produce any significant changes in the mitochondrial morphology of macrophages. The footprint is defined as the total area covered by signal after separated from the background in the image. It is simply the number of pixels in the image containing information of the mitochondria multiplied by the area of a pixel if the calibration is carried out. Similarly, Fig. 6 (b) is plotted for 33, 25 and 38 cells for control, LPS and TNF-*α* challenge trophoblasts, respectively. For trophoblasts, LPS does not produce any kind of morphological changes in mitochondria as can be seen from Fig. 6 (b) while TNF-*α* changes the count of mitochondria after inflammation. The results taken together from cell membrane and mitochondria suggests that the TNF-*α* does not produce directly a detectable level of inflammation in macrophages while LPS changes the morphology at cellular and sub-cellular level. Similarly, LPS does not produce directly detectable level of inflammation in trophoblasts while TNF-*α* does.

**Fig.6.**
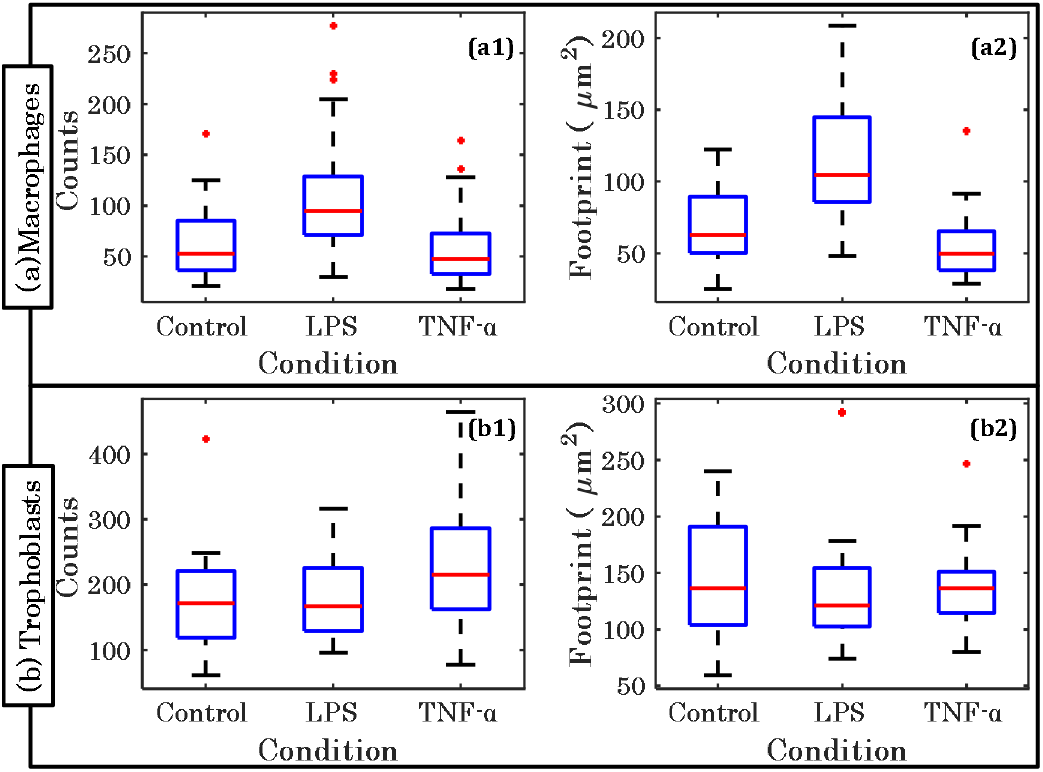
Quantification of the mitochondrial parameters by MiNA image processing tool. (a1), (a2) show the change in mitochondria counts and mitochondrial foot print area, respectively after LPS or TNF-α challenge in macrophages, and (b1), (b2) show the change in mitochondria counts and mitochondrial foot print area, respectively after LPS or TNF-α challenge trophoblasts, respectively after LPS or TNF-α challenge. * shows the outliers data points in dataset.

After confirming the direct inflammatory effects of LPS to macrophage and TNF-α to trophoblasts but not vice-versa, we performed experiments to visualize the rate of morphological changes during inflammation progress. This study also provides the information on initial stage of at inflammation as detected by SIM. The results of SIM show that the changes in the morphology after 2 hr inflammatory agent treatment approximately 20% of the cells are affected at 2 hr time point as shown in Fig. 7. This figure show the change in mitochondrial morphology of the macrophages for different time points of 2 hr, 4 hr and 6 hr of inflammation (Figs. 7 (b-d)) compared to control (Fig. 7 (a)). Figure 7 (e)-(h) show the magnified view of the selected region of the Figs. 7 (a)-(d), respectively. From these images, it is observed that the morphology of the mitochondria is changing even after 2 hours of inflammation compared to the control. It is interesting to note that at 2 hr, NO measurements was not detected in LPS-induced macrophages, as shown in Fig. 1.

**Fig. 7.**
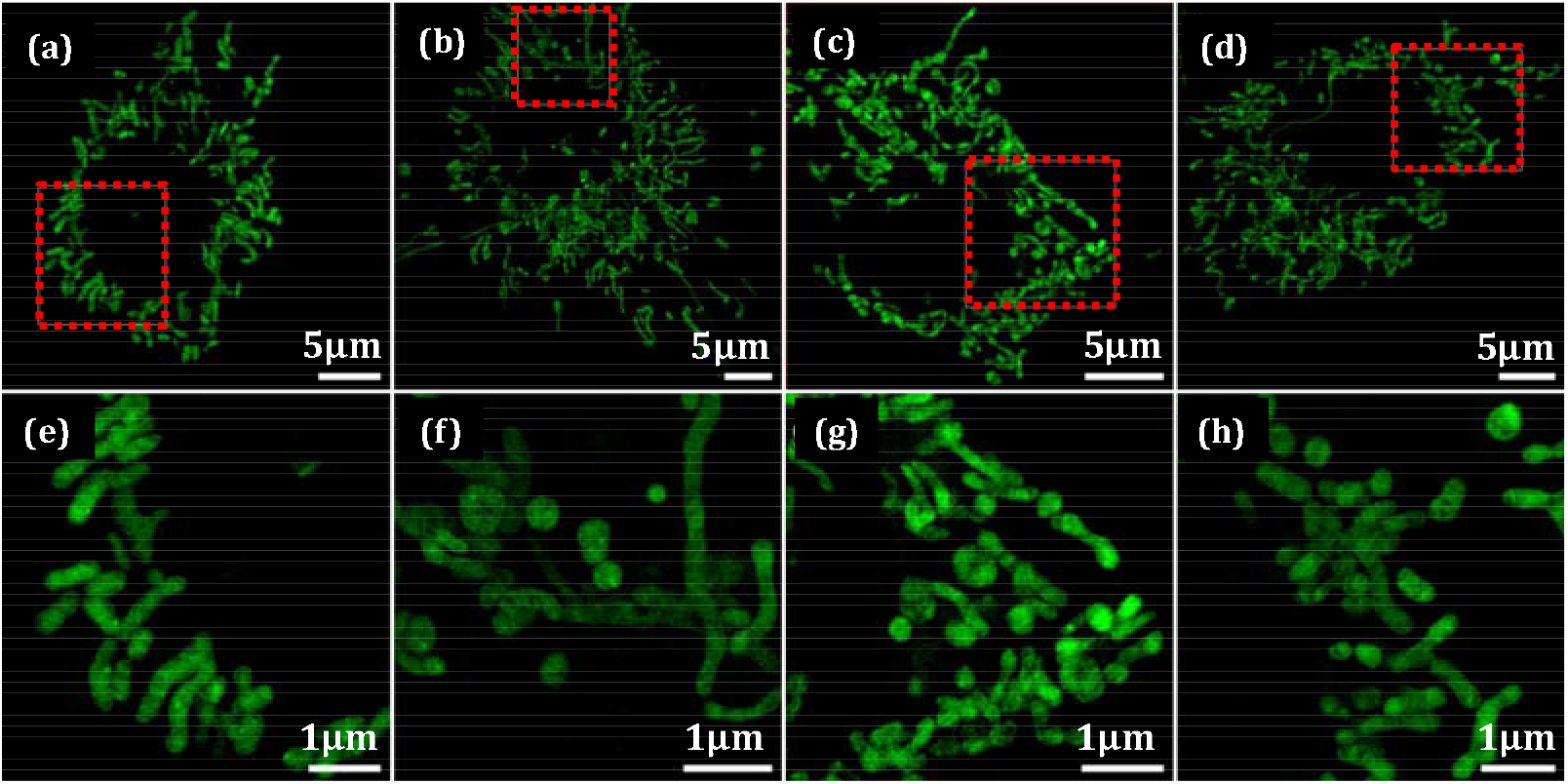
Early time-point imaging of mitochondria of macrophages challenged with LPS. (a) Control; (b), (c) and (d): macrophages challenged with LPS after 2 hr, 4 hr and 6 hr, respectively. (e)-(h) are the magnified image of selected areas of (a), (b), (c) and (d), respectively. Here we can see that the mitochondrial morphology changes after 2 hr compared with control.

### 3.3 Quantitative phase imaging of macrophages and trophoblasts

The QPM of the macrophages and trophoblasts allow us to analyse the optical topography of the specimen and quantify different morphological parameters of the specimen[26, 33]. The quantitative phase information and the details of the morphological changes occurring in the specimen caused by LPS or TNF-*α* on the macrophages and/or trophoblasts could be a standard clinical application. The Linnik based digital holographic microscopy was employed to extract the phase of the specimen. Figure 8 (a)-(c) show the phase images for control, LPS-challenged and TNF-*α*-challenged macrophages, respectively. From Fig. 8 (b), it is observed that LPS exposed macrophages have changed cell morphology. The phase images clearly show that there is decrease in the maximum phase of LPS challenged macrophages as compared to the normal one. The phase image of cell maps the optical path length at each spatial point of the cell, which is a product of the cell thickness and refractive index contrast of the cells and the culture medium. Assuming the refractive index does not change significantly, the net decrease of the maximum phase suggests the thickness of the cell has decreased after LPS challenge. It can also be observed from Fig. 8 (b) that there is an increase in the size of the cell after inflammation as also noticed in SIM images as shown in Fig. 2 (b). Thus, the LPS challenged inflammation on macrophages makes the cell flatter (optically thin) and bigger in area. Contrary, TNF-α does not induce any changes in the phase images of the macrophages as shown in Fig. 8 (c) where the phase values of the TNF-*α*-challenged cells are almost similar to normal one. Also, it does not induce any significant effects on the size of the cells.

**Fig.8.**
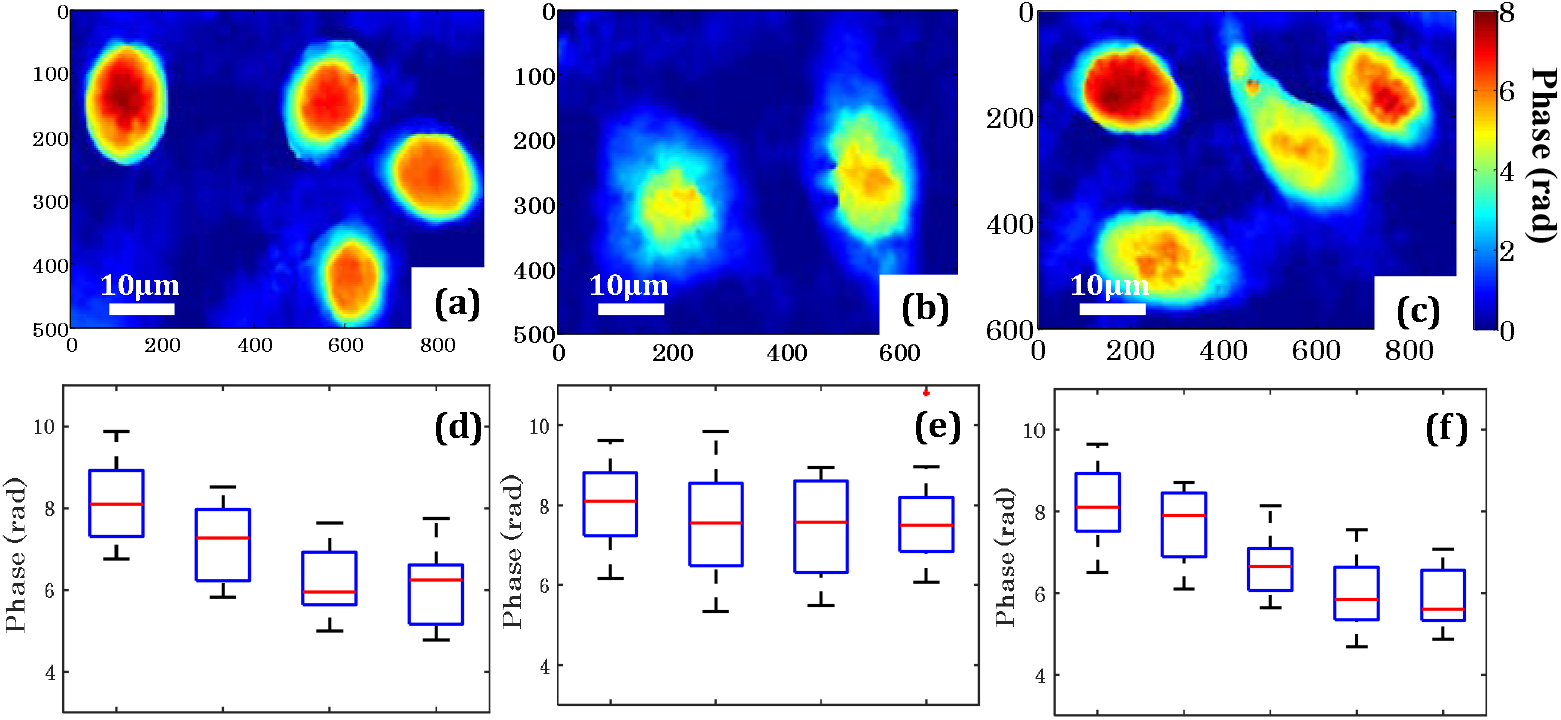
Quantitative phase images of the macrophages and change in the phase map after LPS and TNF α induced inflammation. Figures (a)-(c) show the representative phase images of macrophages for control, LPS-challenged and TNF-α-challenged, respectively. Figures (d) and (e) show LPS and TNF-α concentration dependent change in the maximum phase of macrophages, respectively. Figure (f) shows the time dependent change in the maximum phase of macrophages after challenging with LPS (l μg/ml). * shows the outliers data points in dataset.

**Fig.9.**
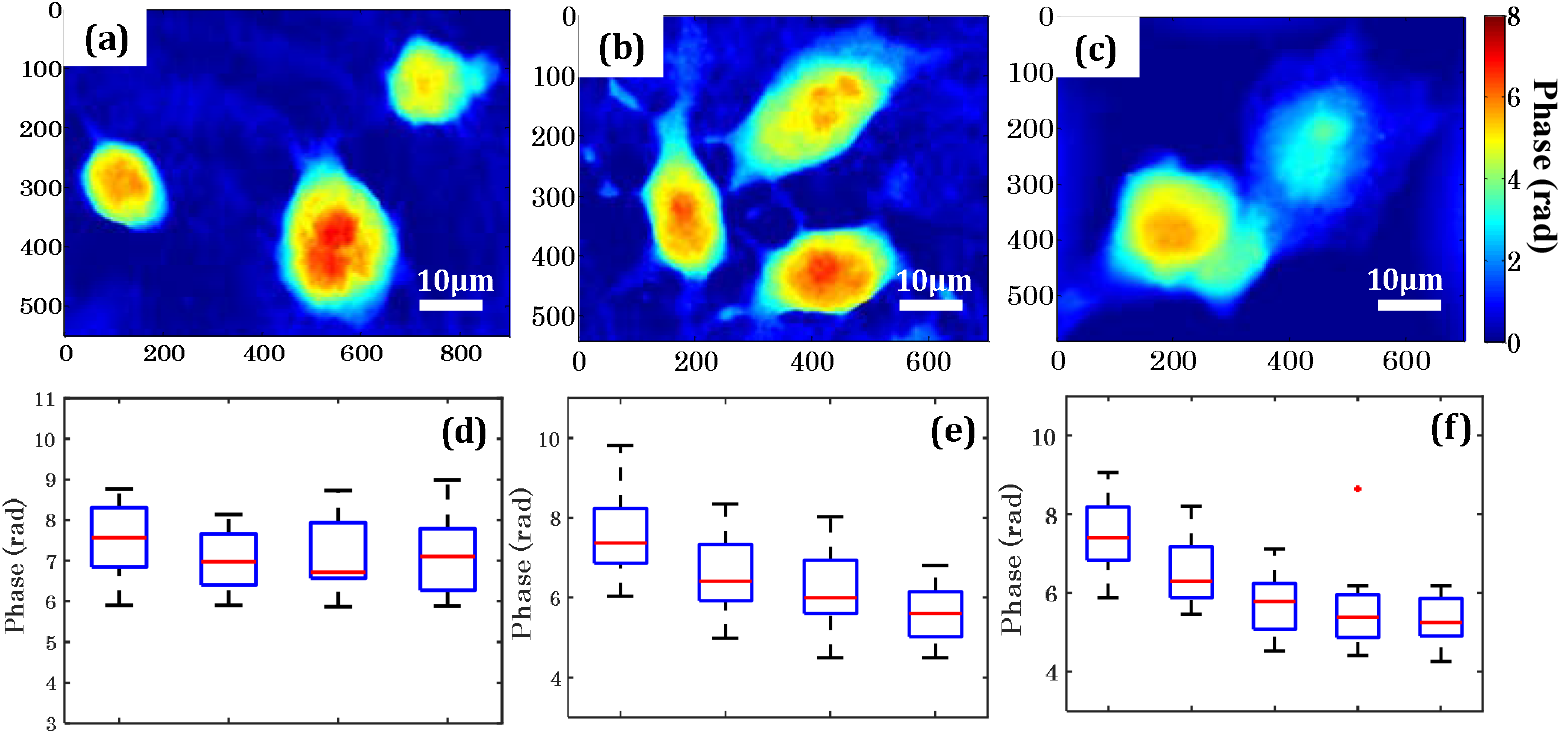
Quantitative phase images of the trophoblasts and change in the phase map after LPS and TNF-α-induced inflammation. Figure (a)-(c) show the representative phase images of trophoblasts for control, LPS challenge and TNF-α challenge respectively. Figure (d) and (e) show LPS and TNF-α concentration dependent change in the maximum phase of trophoblast, respectively and (f) show time dependent change in the maximum phase of trophoblasts after inducing with TNF-α (1000 pg/ml). * shows the outliers data points in dataset.

We have studied the effects of these inflammatory reagents in detail by performing the experiment over different batches and passes of macrophages and trophoblasts and repeated several times (around 200 cells of each case). Figure 8 (d) show the box plots of maximum phase of the macrophages challenged with different concentrations (0.1 μg/ml, 1 μg/ml and 10 μg/ml) of LPS. Results shows that as LPS concentration increased the phase values decrease. There is no change in the maximum phase (optical thickness) of the macrophages observed after challenging different concentrations (10 pg/ml, 100 pg/ml and 1000 pg/ml) of TNF-α as shown in Fig. 8(e). Further, the phase imaging at different time points for a fixed concentration of 1 μg/ml of LPS, shows that the QPM system is capable to detect the minute changes in the morphology just after 2 hr of LPS treatment as shown in Fig. 8(f). This is contrary, under similar condition, measurable quantity of NO production was observed only after 8 hr of LPS-induced inflammation.

Similarly, the effect of various concentration of LPS (0.1 μg/ml, 1 μg/ml and 10 μg/ml) and TNF-*α* (10 pg/ml, 100 pg/ml and 1000 pg/ml) on the morphology of trophoblasts are also studied in detail using QPM. Figure 9 (a)-(c) show the representative phase maps of control, LPS-challenged and TNF-*α*-challenged trophoblasts, respectively. It is observed from Fig. 9 (b) that LPS does not change the trophoblasts morphology while TNF-*α* induces reduction of the maximum phase of trophoblasts as shown in Fig. 9(c). An increment in the cell size is observed after treatment with TNF-*α* which is otherwise not visible in LPS-challenged trophoblasts. Further, the concentration dependent variation in the phase of trophoblast are also quantified as a function of concentration shown in Fig. 9 (d), (e). By the result of phase image and quantitative data, it can be concluded that LPS do not show direct effect on trophoblast while TNF-*α* affects trophoblasts by causing on reduction in the maximum phase (optical thickness) of the specimen. Figure 9 (f) show the change in the phase of trophoblasts at different time points after inducing the 1000 pg/ml concentration of TNF-*α*. The results show that the QPM system is also capable of detecting the early stage changes in the morphology of the trophoblasts at initial time point of 2 hr.

QPM provides the label-free imaging for the quantification of the effect of various inflammatory reagents as well as it is capable to detect the changes in the morphology of the cells after inflammation. These results having a good agreement with the outcomes of the SIM. QPM results contain the information about the change in the optical thickness distribution of the cells as well as various morphological parameters during inflammation. The quantification of these parameters can be helpful for the classification control and inflamed cells in very early stage. Therefore, exploiting machine learning, a support vector machine (SVM)[37, 38] based classifier has been developed and trained with the outcomes of QPM results for the classification of the normal and inflamed cells.

### 3.4 Feature extraction and classification of control and inflamed cells

Application of machine learning tools to optical microscopy and medical imaging provided the new dimension[37-39]. Single shot phase images acquired using QPM can be used to acquire large set of data without the need of extra-labelling step. This makes QPM an attractive route to explore with machine learning[26, 40]. To this end, a support vector machine (SVM) based classifier was developed for the classification of the inflamed macrophages and trophoblasts as the pathological cells comparing to normal healthy cells. As the focus is to predict early stage of infection, we measured the parameters of the cells with LPS treatment for macrophage and TNF-α treatment for trophoblast after 2 hr. A total number of 200 cells of each condition were used for this study.

Several morphological and texture parameters were derived from the acquired phase images of control and inflamed macrophages or trophoblasts at 2 hr time point. We observed that TNF-α did not induce the cascades of inflammation in macrophages and similarly, LPS does induce any direct cascades of inflammation in trophoblasts. Therefore, for the classification of control vs. inflamed cells, we considered only two cases (a) control vs. LPS-challenged macrophages and (b) control vs. TNF-α-challenged trophoblasts. We have extracted number of morphological parameters such as optical thickness (OT), area (A), volume (V) and A/V ratio as well as number of texture parameters[26] (such as mean, variance, entropy, kurtosis and skewness) of phase distribution over the cell structure. These parameters are extracted in order to measure the deformation in the morphology and phase distribution of the specimen after inflammation. In Table 2 we have listed morphological and texture parameters of the control and inflamed macrophages and trophoblasts for the classification.

**Table 2.** The morphological and texture parameters for controls vs. inflamed macrophages (1 μg/ml concentration of LPS) and trophoblasts (1000 pg/ml concentration of TNF-α) at 2 hr of time point.

| Parameter | Definition | Macrophages |  | Trophoblasts |  |
| --- | --- | --- | --- | --- | --- |
|  |  | Control | 2 hr | Control | 2 hr |
| Max Phase (rad) | $\phi$ | 8.176 | 7.375 | 7.766 | 6.639 |
| OT (μm) | $OT = \phi(x, y) * \lambda / 4\pi$ | 0.408 | 0.367 | 0.388 | 0.331 |
| Area (μm<sup>2</sup>) | $dS = dx dy \sqrt{1 + G_x^2 + G_y^2}$ | 364.554 | 703.315 | 525.53 | 1079.65 |
| Volume(fl) | $V = \iint OT(x, y) dx dy$ | 49.91 | 51.98 | 57.372 | 119.432 |
| A/V (μm<sup>-1</sup>) | $A/V$ | 7.806 | 13.879 | 9.932 | 13.741 |
| Mean | $\frac{1}{N} \sum_{i=1}^N \phi_i$ | 5.188 | 3.46 | 4.065 | 2.935 |
| Variance | $\frac{1}{N} \sum_{i=1}^N (\phi_i - \mu)^2$ | 6.1 | 2.7 | 4.035 | 2.691 |
| Entropy | $-\sum_{i=1}^N p(\phi_i) \log_2 p(\phi_i)$ | 1.154 | 1.822 | 0.95 | 1.712 |
| Kurtosis | $\frac{1}{N} \sum_{i=1}^N \left[ \frac{(\phi_i - \mu)^4}{\sigma^4} \right] - 3$ | 1.855 | 1.693 | 1.87 | 1.836 |
| Skewness | $\frac{1}{N} \sum_{i=1}^N \left[ \frac{(\phi_i - \mu)^3}{\sigma^3} \right]$ | -0.319 | -0.032 | 0.143 | 0.136 |

The SVM algorithm is a binary classifier which can detect the inflammation in macrophages and trophoblasts. The receiver operating characteristic (ROC) curve is calculated for better classification at early time points. More theoretical details about SVM and ROC can be found elsewhere [26, 38, 41]. We have used only non-correlated morphological (max phase, OT and area) and texture (mean, variance, entropy, kurtosis and skewness) parameters as input variables and the actual state of the inflammation as response variable i.e. 0 for control and 1 for inflammation state. For the training of the classifier, 60% of the total 400 cells are used by random selection and rest 40% are used as testing data. The sensitivity, specificity and area under curve (AUC) is calculated to measure the accuracy of the classifier. Figure 8 show the ROC curve for the classification of control vs. inflamed macrophages or trophoblasts. The accuracy for the classification of LPS-challenged macrophages is 99.9% as shown in Fig. 10(a). For TNF-*α*-challenged trophoblasts, the accuracy of the classifier is achieved 98.35% as depicted in Fig. 10(b).

**Fig.10.**
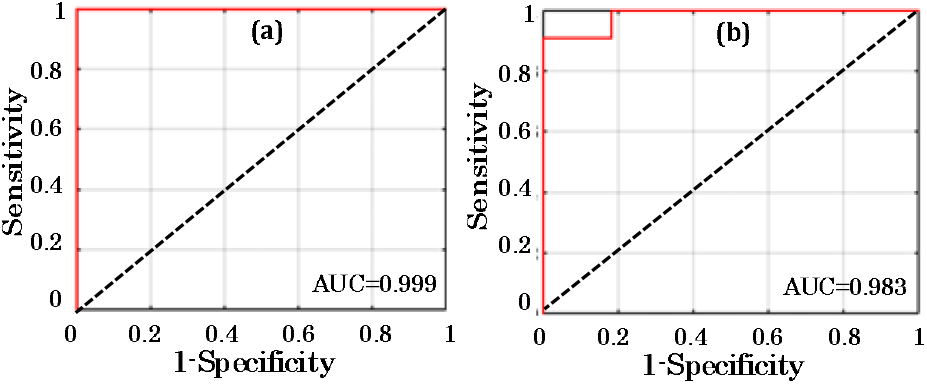
ROC curve for testing dataset for the classification of controls vs. inflamed (a) macrophages or (b) trophoblasts. All the dataset for inflamed cells used for the classification are at 2 hr time point.

## 4. Conclusion

In present study, we have quantified the cellular and sub-cellular morphological and functional changes in macrophages and trophoblasts in response to two inflammatory agents i.e. LPS and TNF-α. We have used super-resolution SIM and label-free QPM to study the effect of these inflammatory reagents on the macrophages and trophoblasts and the results were analysed using SVM-based classifier. It is exhibited that if macrophages or trophoblasts were exposed to similar inflammatory conditions for an equal amount of time, their response to different inflammatory reagents are very different. Multimodal imaging results suggest that the effect of the same inflammatory agent show the different responses to the cellular and sub-cellular morphology of macrophages or trophoblasts. LPS induces significantly large amount of NO production in macrophages which in turn also affects the morphology of mitochondria and the plasma membrane as depicted by SIM images. It also affects either the thickness of the cell or the cellular content (RI) as revealed by QPM results. Contrarily, using high-resolution SIM microscopy and QPM, it was found that more than 60% of the macrophages undergo morphological changes following 24 hr LPS-challenge. However, another inflammatory agent such as TNF-α did not show any direct effect to macrophages.

Particularly for trophoblasts, no significant amount of NO produced even after inflammation induced by both inflammatory agents (LPS or TNF-*α*) because of lacking iNOS presence, however, the sub-cellular changes can be easily detected and quantified using SIM and QPM. Through SIM and QPM, it was also found that the effect on the morphology changes of macrophages and trophoblasts can be observed following just 2 h of incubation with LPS or TNF-*α*. respectively, whereas the NO production in LPS-induced microphage could not be detected at 2 hr. SIM results suggest that the damage in the plasma membrane was more prominent than in mitochondria at early time-points compared with control samples, and the size of cells started to increase following only 2 hr of incubation. SIM results also shows that TNF-*α* is not changing the mitochondrial morphology in trophoblasts. QPM results show that the morphology of LPS-challenged macrophages or TNF-*α*-challenged trophoblasts are drastically changing. The effect of different concentration of LPS or TNF-*α* on the morphology of macrophages or trophoblasts are also quantified with the phase imaging of the cells.

Super-resolution SIM and QPM techniques can be applied directly to live-cell, a few cells are sufficient to detect and distinguish the pathological conditions, multiple cell types can be analysed at the same time, and the state of inflammation can be detected quantitatively in the early stage. It is early to apply these basic data clinically at this stage, however, multi-modal advanced microscopy techniques coupled with machine learning could be the useful tools to evaluate sub-cellular mechanisms associated with inflammation-mediated pregnancy complications with immense potential to explore this untouched field further for clinical applications. In particular, we believe that QPM being label-free and high-speed imaging method when coupled with machine learning can penetrate the clinical imaging demands. Here, we demonstrated that a simple SVM based classifier can achieves an accuracy of 99.9% for LPS-challenged macrophages and 98.3% for TNF-*α*-challenged trophoblasts.

## Acknowledgements

BSA acknowledge the support from the Norwegian Agency for International Cooperation and Quality Enhancement in Higher Education DIKU-Norway (Project number INCP-2014/10024) and the Tematiske Satsinger funding program by UiT The Arctic University of Norway. B.S.A acknowledges the funding from the Research Council of Norway, (project # NANO 2021 – 288565) and (project # BIOTEK 2021 – 285571).

## Author Contributions

RS and VD lead the project and prepared the cells, analysed the results and performed the imaging experiments. RS and DW performed SIM imaging, RS and PB performed NO measurements, VD and AA performed the QPM experiments, VD and AB worked on the machine learning data. AA, VD, AB, build and calibrated the QPM system, DSM and BSA also contributed in the QPM. RS, VD, DW and PB assisted in cell culture and cell handling. PB, GA, DW and RS designed the biological experiments and VD, AA, AB, BSA and DSM designed the QPM experiments. RS and VD mainly wrote the paper and all authors contributed to or commented on the manuscript. BSA, PB and GA conceived the project idea. BSA and PB supervised overall work and BSA provided the funding for the project.

